# Metabolic regulation inferred from Jacobian and Hessian matrices of metabolic functions

**DOI:** 10.1101/2021.10.05.463227

**Authors:** Thomas Nägele

**Author notes:** **Correspondence:**, T.: 0049-89-2180-74660.

## Abstract

Quantitative analysis of experimental metabolic data is frequently challenged by non-intuitive, complex patterns which emerge from regulatory networks. Quantitative output of metabolic regulation can be summarised by metabolic functions which comprise information about dynamics of metabolite concentrations. They reflect the sum of biochemical reactions which affect a metabolite concentration. Derivatives of metabolic functions provide essential information about system dynamics. The Jacobian matrix of a reaction network summarises first-order partial derivatives of metabolic functions with respect to metabolite concentrations while Hessian matrices summarise second-order partial derivatives. Here, a simple model of invertase-driven sucrose hydrolysis is simulated and both Jacobian and Hessian matrices of metabolic functions are derived for quantitative analysis of kinetic regulation of sucrose metabolism. Based on previous experimental observations, metabolite dynamics are quantitatively explained in context of underlying metabolic functions. Their potential regulatory role during plant cold acclimation is derived from Jacobian and Hessian matrices.

## Introduction

The quantitative study of biochemical reaction networks represents an interdisciplinary research area of (bio)chemistry, physics and mathematics. Enzymes catalyse chemical reactions under physiologically relevant conditions. Enzyme activity directly depends on temperature, pH, ion strength and redox potential of a cell or compartment showing characteristic optima (Arcus &Mulholland, 2020; Bisswanger, 2017). In addition, enzyme activity in cellular systems is affected and regulated by diverse biochemical effectors, e.g., comprising other proteins and metabolites (Atkinson, 1969; Chen et al., 2021). As a result, cellular enzyme activity represents a variable of biochemical networks which is shaped by a large parameter space challenging experimental, but also theoretical, analysis. Enzyme kinetic models mathematically describe enzymatic reaction rates as a function of one or more parameters and variables. In general, biochemical kinetics is based on the mass action law assuming the reaction rate to be proportional to the probability of reactant collision (Waage &Gulberg, 1864, 1986). This probability is proportional (i) to the concentration of reactants, and (ii) to the number of molecules of a species which participate in a reaction, i.e., to the power of molecularity. The rate *v* of a reaction following the mass action law with molecularities *m*_*i*_ and *m*_*j*_ of substrates *S*_*i*_ and products *P*_*j*_, respectively, is described by the rate equation (Eq. 1):

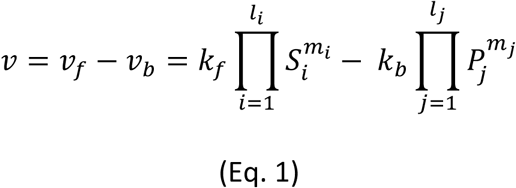

Here, *k*_*f*_ and *k*_*b*_ represent the rate constants, i.e., proportionality factors, for the forward (*k*_*f*_) and backward (*k*_*b*_) reaction. Concentration dynamics of involved substrate and product molecules can be described by differential equations (DEs). If concentration dynamics are considered (only) over time, ordinary differential equations (ODEs) are applied while partial differential equations account for more than one independent variable, e.g., time and space. Concentration dynamics over time of an arbitrary enzyme reaction which interconverts a substrate molecule S_1_ into a product molecule P_1_ are then described by the following ODE (Eq. 2).

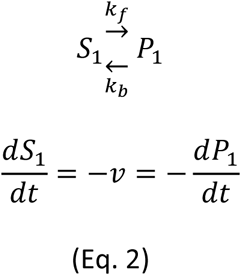

### Deriving a Jacobian matrix to study invertase-catalysed sucrose hydrolysis

Introducing the reversible formation of an enzyme-substrate complex (E + S → ES), a release of product P from ES (ES → E + P) (Brown, 1902), and the simplifying assumption that formation of ES is much faster than its decomposition into E and P finally yields the Henri-Michaelis-Menten kinetics (Henri, 1902, 1903; Michaelis &Menten, 1913) (Eq. 3).

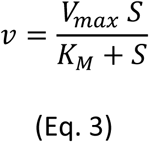

The parameter V_max_ represents the maximal velocity which is reached when the enzyme is completely saturated with substrate. The parameter K_M_ represents the Michaelis constant which equals the substrate concentration at V_max_/2. A more detailed derivation and an explanation of kinetics in context of reaction thermodynamics has been provided earlier, see e.g. (Klipp et al., 2016). Due to its capability to accurately describe and quantify mechanisms of enzyme catalysis and regulation, the Michaelis-Menten equation is crucial for biochemical understanding (Cornish-Bowden, 2015). It was derived based on experimental observations of sucrose hydrolysis, catalysed by invertase enzymes (Brown, 1902; Michaelis &Menten, 1913). Within this reaction, the glycosidic bond of sucrose is hydrolysed, and glucose and fructose are released (Eq. 4).

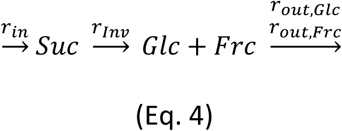

In this kinetic model, *r*_*in*_ represents the rate of sucrose biosynthesis, *r*_*inv*_ the rate of invertase-driven hydrolysis and *r*_*out,Glc*_ and *r*_*out,Frc*_ hexose consuming processes, e.g., phosphorylation by hexokinase enzymes. The corresponding ODE model of this reaction system describes sucrose, fructose and glucose dynamics by the sum of in- and effluxes (Eqs. 5-7).

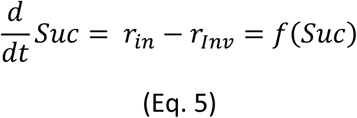

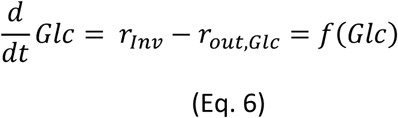

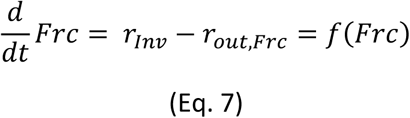

The right side of the ODEs, i.e., the sum of reactions, is summarised by metabolic functions ***f*** and their integration yields the time course of metabolite concentrations.

Invertases play a central role in diverse processes of plant metabolism, development and response to environmental stress (Koch, 2004; Ruan, 2014; Vu et al., 2020; Weiszmann et al., 2018; Xiang et al., 2011). Plant invertases occur in different isoforms with different compartmental localisation and biochemical properties (Sturm, 1996; Tymowska-Lalanne &Kreis, 1998). Both plant vacuolar and extracellular invertases possess an acidic pH optimum between 4.5 and 5.0 while cytosolic invertase has a neutral pH optimum between 7.0 and 7.8 (Sturm, 1999). Acid and neutral invertases hydrolyse sucrose with a K_M_ in a low-millimolar range (Sturm, 1999; Unger et al., 1992). Invertases are product inhibited, with glucose acting as a non-competitive inhibitor and fructose as a competitive inhibitor (Sturm, 1999).

Due to its central role in sugar metabolism, a quantitative understanding of invertase kinetics is crucial for analysis and interpretation of photosynthesis and plant carbohydrate metabolism. Also, sugar signalling, plant fertility and fitness are significantly affected and regulated by invertases which emphasises even more their essential role in plants (Wan et al., 2018). Yet, as soon as enzyme kinetics recorded *in vitro* are applied to explain *in vivo* metabolic regulation and to simulate metabolic fluxes, further information about involved metabolite pools, compartments and enzyme regulation is needed in order to estimate dynamics of substrate (*here*: sucrose) and product (*here*: hexose) concentrations. If metabolite concentrations and kinetic parameters are known, the system of ODEs which describes dynamics of sucrose and hexose concentrations (Eqs. 5-7) can be numerically solved, i.e., numerically integrated. Thus, to estimate the impact of product inhibition of invertases, dynamics of hexoses need to be estimated which depend on invertase activity (input) and output reactions which consume or interconvert glucose and fructose (here: *r*_*out,Glc*_ and *r*_*out,Frc*_). A physiologically important output reaction, which interconverts hexoses, is their phosphorylation, catalysed by hexokinases (Granot et al., 2014). To prevent the depletion of sucrose, an input function needs to be defined which supplies the system with sucrose molecules (*r*_*in*_). In plant metabolism, sucrose phosphate synthase (SPS) catalyses and regulates sucrose biosynthesis in the cytosol together with fructose-1,6-bisphosphatase (FBPase) (Doehlert &Huber, 1983; Stitt et al., 1983; Volkert et al., 2014). Here, the input reaction rate *r*_*in*_ was defined to be constant without explicitly modelling SPS or FBPase kinetics (Eq. 8). Invertase reaction and hexose output followed Michaelis-Menten kinetics considering product inhibition of invertase (Eq. 9) while output was assumed not to be inhibited (Eqs. 10, 11).

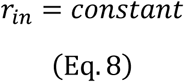

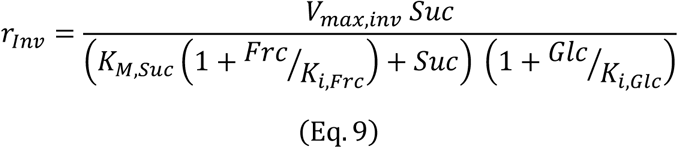

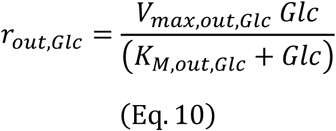

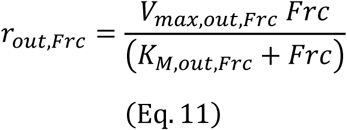

Parameters *V*_*max*,…_ are the maximum velocities of enzyme reactions, i.e., reaction rates under substrate saturation. Michaelis constants K_M,…_ are affinities of enzymes for their substrates and represent the metabolite concentration at r = V_max_/2. The inhibitory constants K_i,…_ indicate inhibitor concentrations which are needed to reduce enzyme velocity to half maximum. To study complex system properties like stabilization after perturbation, biochemical systems are typically linearised and considered near a steady state. Yet, due to strong external and internal dynamics, plant metabolism can hardly be described by steady state assumptions, which is d**M(t)**/dt = **0** where **M(t)** represents a vector of metabolite concentrations and **0** is the zero vector. In the given example of sucrose hydrolysis within plant cells (Eq. 4), both substrate and product molecules may show significant dynamics within a diurnal cycle (Sulpice et al., 2014). Although this clearly constrains the interpretation of findings made by assumptions of a steady state, it simplifies theoretical and computational analysis of metabolic systems. Additionally, analytical solutions of enzymatic reaction systems can be obtained using the assumption that velocity of an enzymatic reaction linearly depends on substrate concentrations (Heinrich &Rapoport, 1974). In contrast, solving nonlinear metabolic systems analytically is hardly possible. Taylor expansion of temporal changes of deviations from a steady state leads to the Jacobian matrix of a reaction system (Klipp et al., 2016). In context of metabolic networks, the Jacobian matrix describes elasticities of metabolic functions towards dynamics of metabolite concentrations. For the given reaction system (Eq. 4), the Jacobian matrix reads (Eq. 12):

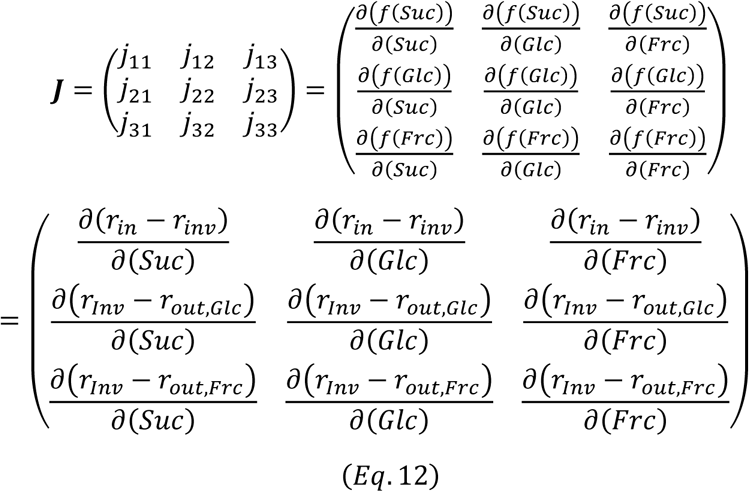

with diagonal entries (Eqs. 13-15),

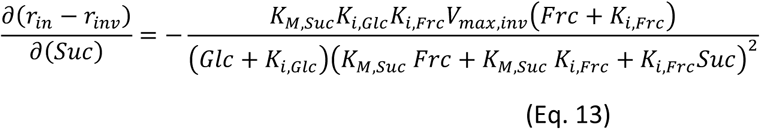

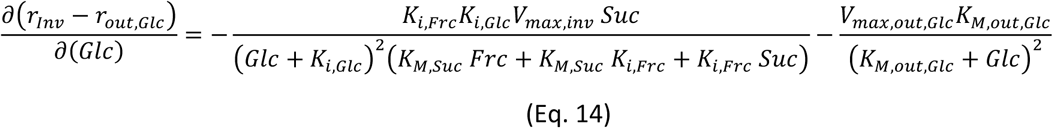

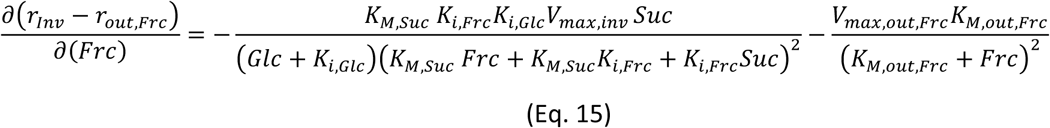

In words, the diagonal entries represent elasticities of functions of metabolites with regard to dynamics in their own concentration. This reflects that metabolites are affecting their own concentration dynamics by participating as substrates and/or products in enzymatic reactions. Non-diagonal entries of ***J*** reflect any regulatory effects of metabolites on all other metabolic functions within a reaction system (equations of derivatives are not explicitly shown). Further, the equations show that, depending on the degree of substrate saturation of an enzyme, dynamics of substrate concentrations have a differential effect on the Jacobian trajectories. For example, if enzymes, catalysing the *r*_*out,Glc*_ and *r*_*out,Frc*_ reactions, are fully saturated, further increase of substrate concentration (here: Glc and Frc) results in square decrease towards zero of the second term in Eqs. 14 and 15. Hence, under such conditions where reaction products of invertases strongly accumulate, hexose consuming reactions have a lower impact on hexose dynamics than under non-saturated conditions. At the same time, (strong) hexose accumulation results in invertase inhibition which limits invertase-induced sucrose dynamics because (see Eq. 13):

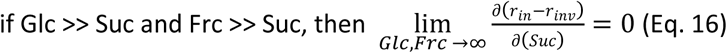

These considerations are simplified because also enzyme parameters need to be considered in order to determine the Jacobian entries as well as their dynamics under different ratios of substrate-product concentrations. While K_M_ and K_i_ represent characteristic and constant enzyme parameters, V_max_ might differ significantly between time points and conditions under which (plant) metabolism is analysed. This might be due to a changing total enzyme amount or a changing temperature which affects reaction constants as described by the Arrhenius equation (Arrhenius, 1889). Altogether, this establishes a multi-parameter space for the estimation of a biochemical Jacobian matrix. Its estimation from experimental data is challenged by the need for kinetic parameters which are sometimes not available or need to be acquired within laborious, difficult and error-prone experiments.

To overcome experimental limitations by forward kinetic experiments, inverse estimation of ***J*** has been suggested based on covariance information of metabolite concentrations (Steuer et al., 2003). Here, fluctuation terms at a metabolic steady state are estimated from (co)variance information contained in metabolomics data. Applying this approach, regulatory hubs of metabolic networks can be identified without measuring enzyme kinetic parameters (Nägele et al., 2014; Sun &Weckwerth, 2012; Weckwerth, 2019; Wilson et al., 2020). However, experimental validation of predicted effects on enzymatic regulation is essential for such inverse estimations. Two main reasons why such inverse approximation might fail to correctly predict metabolic regulation are (i) strong deviation from steady state assumption due to significant internal and/or external perturbations, and (ii) misinterpretation of reasons for metabolite (co)variance, e.g., technical variance. Despite all experimental complications, a combined approach of inverse Jacobian matrix estimation and experimental validation might support the analysis and understanding of complex metabolic network regulation (Wilson et al., 2020).

In summary, calculation and estimation of Jacobian matrices is a crucial element of system theory, and its application to biochemical networks seems mandatory to unravel complex regulatory principles. Entries of the Jacobian matrix are first-order partial derivatives of variable functions. As explained before, in a metabolic network context, it provides information about the effect of dynamics of one metabolite concentration on a specific metabolic function. Also, evaluation of eigenvalues of ***J*** indicates stability or instability of a metabolic system which provides important information about system behaviour at a considered steady state (Fürtauer &Nägele, 2016; Grimbs et al., 2007; Reznik &Segrè, 2010). In the following paragraph, the Hessian matrix is introduced in context of metabolic regulation which comprises second-order partial derivatives. It is discussed in context of the provided example of invertase-catalysed sucrose cleavage.

### Applying the Hessian matrix to study substrate-product-interactions on metabolic functions

The Hessian matrix of a function f(x_1_, x_2_, x_3_, …, x_n_) with n variables contains its second-order partial derivatives. Following Schwarz’ theorem (also: Clairaut’s theorem), it is an *n x n* symmetric matrix, i.e., the second-order partial derivatives satisfy identity regarding the order of differentiation (Eq. 17).

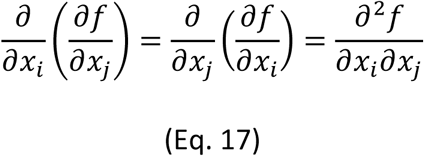

Considering metabolite concentrations as variables within metabolic functions of the given invertase reaction system (Eq. 4), the Hessian matrices of *f(Suc)* calculates as follows (Eq. 18):

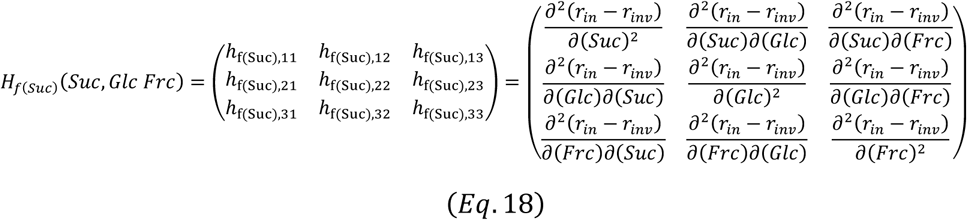

Accordingly, Hessian matrices of metabolic functions of both reaction products, glucose and fructose, comprise second-order partial derivates of *f(Glc)* and *f(Frc)*, respectively (Eqs. 19 and 20).

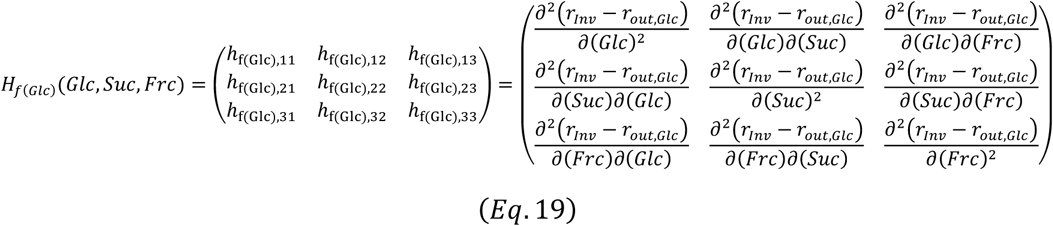

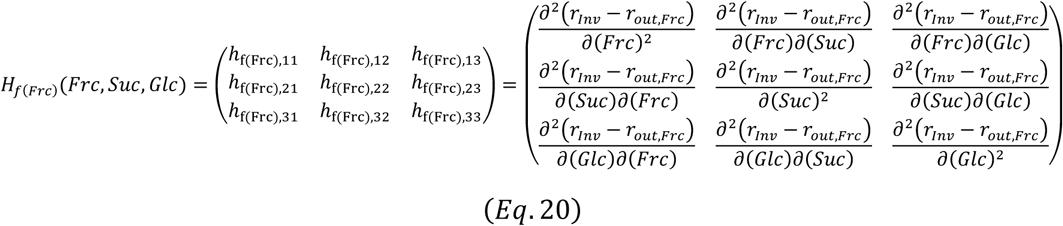

The Hessian matrix provides information about simultaneous effects of a combination of concentration dynamics of substrates, products or other metabolic effectors on a metabolic function. For example, in an invertase reaction (Eq. 4), entries 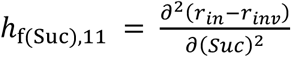, 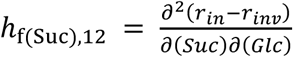 and 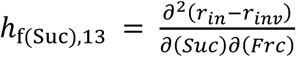 provide information about how the metabolic function of sucrose depends on dynamics of (i) sucrose (*h*_*f(Suc),11*_), (ii) sucrose and glucose (*h*_*f(Suc),12*_), and (iii) sucrose and fructose concentrations (*h*_*f(Suc),13*_), respectively (Eqs. 21-23).

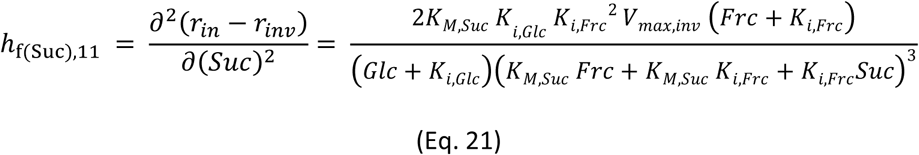

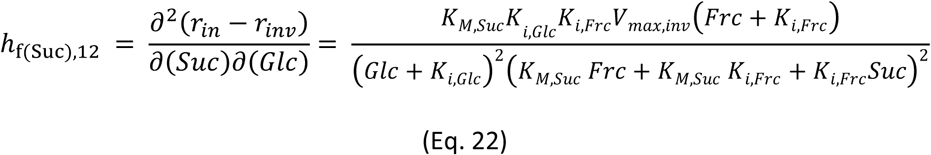

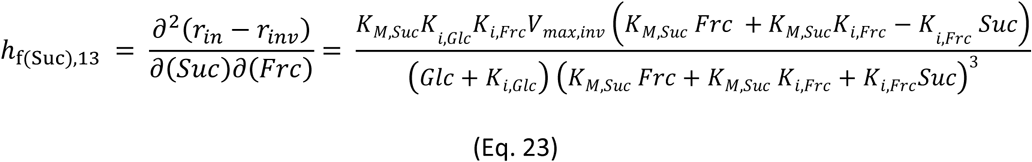

Competitive (Frc) and non-competitive inhibitors (Glc) differentially shape and affect the metabolic function of the reaction substrate (Suc) which is expressed in Hessian terms by different positions and exponents in nominators and denominators (Eqs. 21-23). While differential regulatory impact of different types of inhibitors on enzyme kinetics becomes evident from experimental enzyme kinetic analysis using purified enzymes, its interpretation in context of metabolic functions remains difficult. In this context, Jacobian and Hessian matrices provide important insight because they describe dynamics of metabolic functions with respect to a certain variable, e.g., a metabolic inhibitor. Further, due to plasticity of metabolism, metabolite concentrations may vary significantly under similar environmental conditions and without stress exposure. For example, sucrose and hexoses may accumulate significantly, and even double in amount, during the light period of a diurnal cycle (Brauner et al., 2014; Seydel et al., 2021; Sulpice et al., 2014). Such strong dynamics of reaction product and substrate concentrations aggravate the quantitative analysis of metabolic regulation due to their non-linear impact on enzymatic rates. It follows that instead of analysing one (single) snapshot, a broad range of physiologically relevant metabolite concentrations and/or enzyme parameters needs to be analysed in order to cope with metabolic plasticity. Here, an example for such an analysis is provided applying a kinetic parameter set of invertase reactions (Table I), which has previously been determined in *Arabidopsis thaliana* under ambient (22°C) and low (4°C) temperature (Kitashova et al., 2021).

**Table I.** Carbon uptake rates and kinetic parameters of invertase catalysed sucrose hydrolysis in *Arabidopsis thaliana*, accession Col-0, at 22°C and 4°C. Data source: (Kitashova et al., 2021).

| Kinetic Parameter | Absolute value | Dimension |
| --- | --- | --- |
| $r_{in,22^{\circ}\text{C}}$ | 8 | $[\mu\text{mol Suc h}^{-1} \text{gFW}^{-1}]$ |
| $r_{in,4^{\circ}\text{C}}$ | 0.75 | $[\mu\text{mol Suc h}^{-1} \text{gFW}^{-1}]$ |
| $V_{\text{max,inv},22^{\circ}\text{C}}$ | 35.2 | $[\mu\text{mol Suc h}^{-1} \text{gFW}^{-1}]$ |
| $V_{\text{max,inv},4^{\circ}\text{C}}$ | 2 | $[\mu\text{mol Suc h}^{-1} \text{gFW}^{-1}]$ |
| $K_{M,\text{Suc}}$ | 12 | $[\mu\text{mol Suc gFW}^{-1}]$ |
| $K_{i,\text{Frc}}$ | 0.23 | $[\mu\text{mol Frc gFW}^{-1}]$ |
| $K_{i,\text{Glc}}$ | 0.12 | $[\mu\text{mol Glc gFW}^{-1}]$ |

Reaction rates of invertase enzymes, *r*_*inv*_, were calculated across different combinations of physiologically relevant sucrose and hexose concentrations to determine the metabolic function of sucrose, i.e., *f(Suc) = r*_*in*_ *-r*_*inv*_. Simulation results of different sucrose concentrations were plotted against glucose and fructose concentrations (Figure 1). Thus, each shown plane in the figure corresponds to solutions of ***f(Suc), J*** and ***H*** for one sucrose concentration (a detailed definition of concentrations is provided in the figure legend). Although sucrose concentrations used for 4°C simulations were up to 8-fold higher than under 22°C, resulting absolute values and dynamics of ***f(Suc)*** were significantly lower than under 22°C (Fig. 1 a and b). Reduced absolute values were due to a decreased input rate *r*_*in,4°C*_ (based on experimental findings). As expected, under conditions of low product concentration, ***f(Suc)*** became minimal under both temperatures due to increased rates of sucrose cleavage (Fig. 1 a, b). However, reduced dynamics of ***f(Suc)*** was due to increased hexose concentrations (inhibitors) and a reduced *V*_*max*_ of invertase (see Table I). As a result, also the dynamic range of ***J*** and ***H*** decreased across all simulated scenarios by several orders of magnitude (10^−1^ → 10^−4^/10^−5^; Fig. 1 c-f). A main low temperature effect became visible in entries of Jacobian matrices which was a reduced degree of overlap between 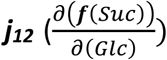 and 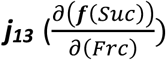 (Fig. 1 c, d). Both terms describe changes of ***f(Suc)*** induced by (slight) changes of glucose and fructose concentrations, respectively. At 22°C, high glucose concentrations (∼2.5-3 µmol gFW^-1^) minimise ***j***_***13***_ and, with this, also the regulatory effect of fructose dynamics on ***f(Suc)*** (see Fig. 1 c). At 4°C, high glucose concentrations (∼14-15 µmol gFW^-1^) also lead to minimal values of ***j***_***13***_, which were, however, still significantly higher than ***j***_***12***_ (see Fig. 1 d; ANOVA, p<0.001). This discrepancy became also visible in the curvature of ***f(Suc)***, i.e., in the Hessian matrix (Fig. 1 e, f).

**Figure 1.**
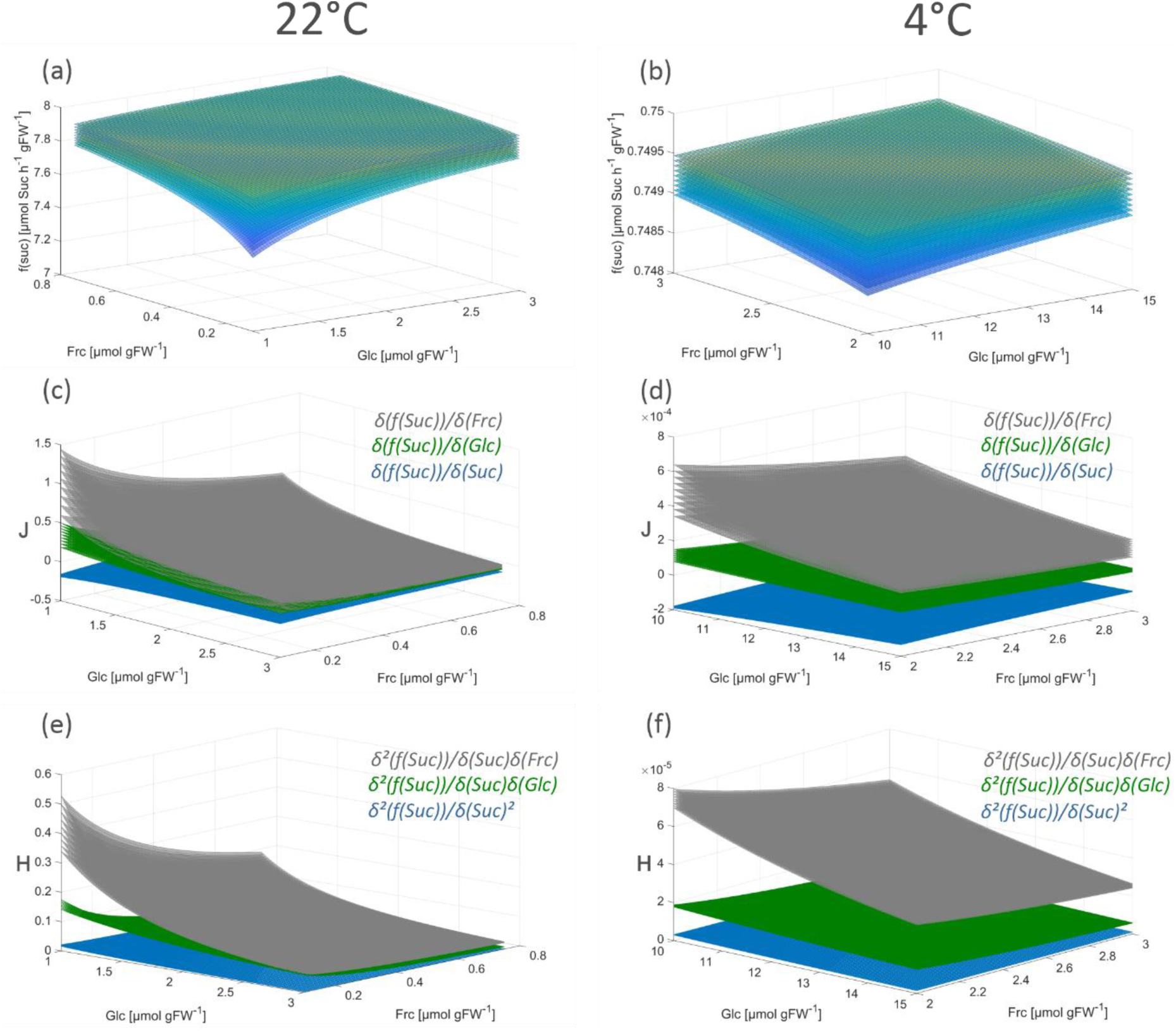
Dynamics of *f(Suc)* under ambient and low temperature. **(a)** *f(Suc)* at 22°C under variable concentrations of fructose (x-axis), glucose (y-axis) and sucrose (planes). **(b)** *f(Suc)* at 4°C under variable concentrations of fructose (x-axis), glucose (y-axis) and sucrose (planes). **(c)** Jacobian matrix entries of *f(Suc)* at 22°C under variable concentrations of glucose (x-axis), fructose (y-axis) and sucrose (planes; see Eq. 12; ***j***_***11***_: blue; ***j***_***12***_: green; ***j***_***13***_: grey). (**d**) Jacobian matrix entries of *f(Suc)* at 4°C under variable concentrations of glucose (x-axis), fructose (y-axis) and sucrose (planes). See also Eq. 12; ***J***_***11***_: blue; ***J***_***12***_: green; ***J***_***13***_: grey. **(e)** Hessian matrix entries of *f(Suc)* at 22°C under variable concentrations of glucose (x-axis), fructose (y-axis) and sucrose (planes), see also Eq. 18; ***h***_***f(Suc,11)***_: blue; ***h***_***f(Suc,12)***_: green; ***h***_***f(Suc,13)***_: grey). **(f)** Hessian matrix entries of *f(Suc)* at 4°C under variable concentrations of glucose (x-axis), fructose (y-axis) and sucrose (planes), see also Eq. 18; ***h***_***f(Suc,11)***_: blue; ***h***_***f(Suc,12)***_: green; ***h***_***f(Suc,13)***_: grey). Each plane corresponds to a sucrose concentration which was varied between 1-3 µmol gFW^-1^ and 4-8 µmol gFW^-1^ for simulations at 22°C and 4°C, respectively.

These observations suggest that, under ambient conditions and (increased) glucose concentrations, it is ***j***_***12***_ ≈ ***j***_***13***_, i.e., 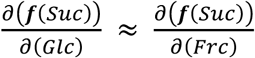, and ***h***_**f(Suc)**,**12**_ ≈ ***h***_**f(Suc)**,**13**_, i.e., 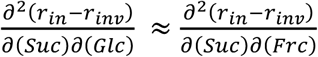. At low temperature, this similarity is not given even under (relatively) high glucose concentrations which might suggest a cold-induced switch of the regulatory role which fructose plays in plant metabolism (Klotke et al., 2004).

## Conclusions

Together with the Jacobian matrix, Hessian matrices are commonly applied to study n-dimensional functions and surfaces, their extrema and their curvature (see e.g. (Basterrechea &Dacorogna, 2014; Ivochkina &Filimonenkova, 2019)). In context of the presented theory for analysis of biochemical metabolic functions, this suggests that metabolism can be summarised by a multi-dimensional function. Finally, Jacobian and Hessian matrices of these functions comprise multi-dimensional quantitative information about dynamics, shape and curvature which might support the identification of underlying regulators of metabolism.

## Acknowledgements

I would like to thank all members of Plant Evolutionary Cell Biology, LMU Munich, for many fruitful seminars and discussions. Special thanks go to Lisa Fürtauer, RWTH Aachen, and AG Weckwerth, University of Vienna, as well as Jakob Weiszmann and Matthias Nagler for constructive advice, support and discussion. Finally, I thank the SFB/TR175 consortium for a supportive research environment and fruitful discussions.

## Financial Support

This work was supported by Deutsche Forschungsgemeinschaft (DFG), grants TR175/D03 and NA 1545/4-1.

## Conflicts of Interest

None

## Notes

### Competing Interest Statement

The authors have declared no competing interest.

